# Microglia cannibalism and efferocytosis leads to shorter lifespans of developmental microglia

**DOI:** 10.1101/2023.03.15.532426

**Authors:** Hannah Gordon, Zachary Schafer, Cody J. Smith

## Abstract

The overproduction of cells and subsequent production of debris is a universal principle of neurodevelopment. Here we show an additional feature of the developing nervous system that causes neural debris – promoted by the sacrificial nature of embryonic microglia that irreversibly become phagocytic after clearing other neural debris. Described as long-lived, microglia colonize the embryonic brain and persist into adulthood. Using transgenic zebrafish to investigate the microglia debris during brain construction, we identified that unlike other neural cell-types that die in developmental stages after they have expanded, necroptosis-dependent microglial debris is prevalent when microglia are expanding in the zebrafish brain. Time-lapse imaging of microglia demonstrates that this debris is cannibalized by other microglia. To investigate features that promote microglia death and cannibalism, we used time-lapse imaging and fate-mapping strategies to track the lifespan of individual developmental microglia. These approaches revealed that instead of embryonic microglia being long-lived cells that completely digest their phagocytic debris, once most developmental microglia in zebrafish become phagocytic they eventually die, including ones that are cannibalistic. These results establish a paradox -- which we tested by increasing neural debris and manipulating phagocytosis -- that once most microglia in the embryo become phagocytic, they die, create debris and then are cannibalized by other microglia, resulting in more phagocytic microglia that are destined to die.

## MAIN TEXT

Programmed cell death and clearance of that debris is a universal principle of nervous system development^1,2^. In vertebrates, this debris is cleared by microglia, which are the resident and professional phagocyte of the brain^3^. Microglia are considered long-lived cells that colonize the embryonic brain from the embryonic yolk-sac^3–7^. This definition is supported by fate-mapping studies that demonstrate yolksac derived cells generate microglia that persist into adulthood and intravital imaging of adult mouse brains that demonstrate low turnover of microglia in the healthy animals^5–8^. Genetic-based lineage-tracing methodologies in zebrafish have also demonstrated embryonic rostral blood island-derived microglia are present in the brain until 15 days post fertilization (dpf)^9,10^. Adult zebrafish microglia are also derived from progenitors that can be labeled in the embryo^10–12^. While these genetic experiments clearly demonstrate that microglia in embryos and adults are produced from embryonic sources, they do not label microglia distinct from their progenitors nor do they allow for sparse labelling that can track the fate of individual microglia. Timelapse imaging of zebrafish microglia has confirmed that the embryonic microglia population is stable over 24-hour periods^13–16^. Conflicting the long-lived nature of individual microglia is evidence of microglia turnover in zebrafish and humans during development, and of microglia death in adult mice^9,17,18^. Thus while the microglia population is stable and long-lived, the lifespan of individual microglia in the embryonic brain remains a concept that is unanswered.

To understand the lifespan of individual microglia, we searched for microglia death during brain construction. Using zebrafish to precisely fate-map individual microglia, we found that once most embryonic microglia become phagocytic, they die. Unique from other cells in development, this death is necroptotic. The scale of this death requires a rapid turnover of microglia when they are expanding in the brain. Unlike the adult brain states when microglia are tiled^19^, our evidence in zebrafish shows that developmental microglia cannibalize microglia debris, resulting in microglia that are destined to die because they are phagocytic. In addition to introducing a new process that produces developmental cell death, this work demonstrates that embryonic microglia lifespan rarely exceeds 3 days when phagocytic debris is prominent.

To explore microglia death in the developing nervous system, we first searched for signatures of microglia death in the developing brain using transgenic zebrafish, *Tg(pu1:Gal4-UAS:GFP)* (hereafter *pu1:GFP*) that use regulatory regions of *pu1* to label microglia populations^15^. To ensure that we could identify death regardless of the mode of death, we first scored the amount of GFP^+^ debris in the brain from 3-5 dpf. In these ages, we scored that GFP^+^ microglia debris was present and increased across days (Figure 1a,b. N=21 animals, p=0.0397 3 dpf vs 5 dpf, Post hoc Tukey Test). We confirmed this debris was derived from microglia and not from other *pu1*^*+*^ macrophages that may enter the embryonic brain by demonstrating they also label with the zebrafish microglia-specific antibody 4C4 (Figure 1c)^20,21^. 4C4 does label a subset of macrophage but they are localized outside the brain region in wildtype animals. To complement the 4C4 labeling, we also used timelapse imaging to visualize death in *Tg(pu1:GFP)* animals at 4 dpf and then scored cells ramified morphology that is typical of microglia. The projection length and number of projections between microglia that were fated to die compared to 4C4^+^ microglia were indistinguishable (Figure 1d,e,f, S1a), consistent with their microglia identity.

**Figure 1.**
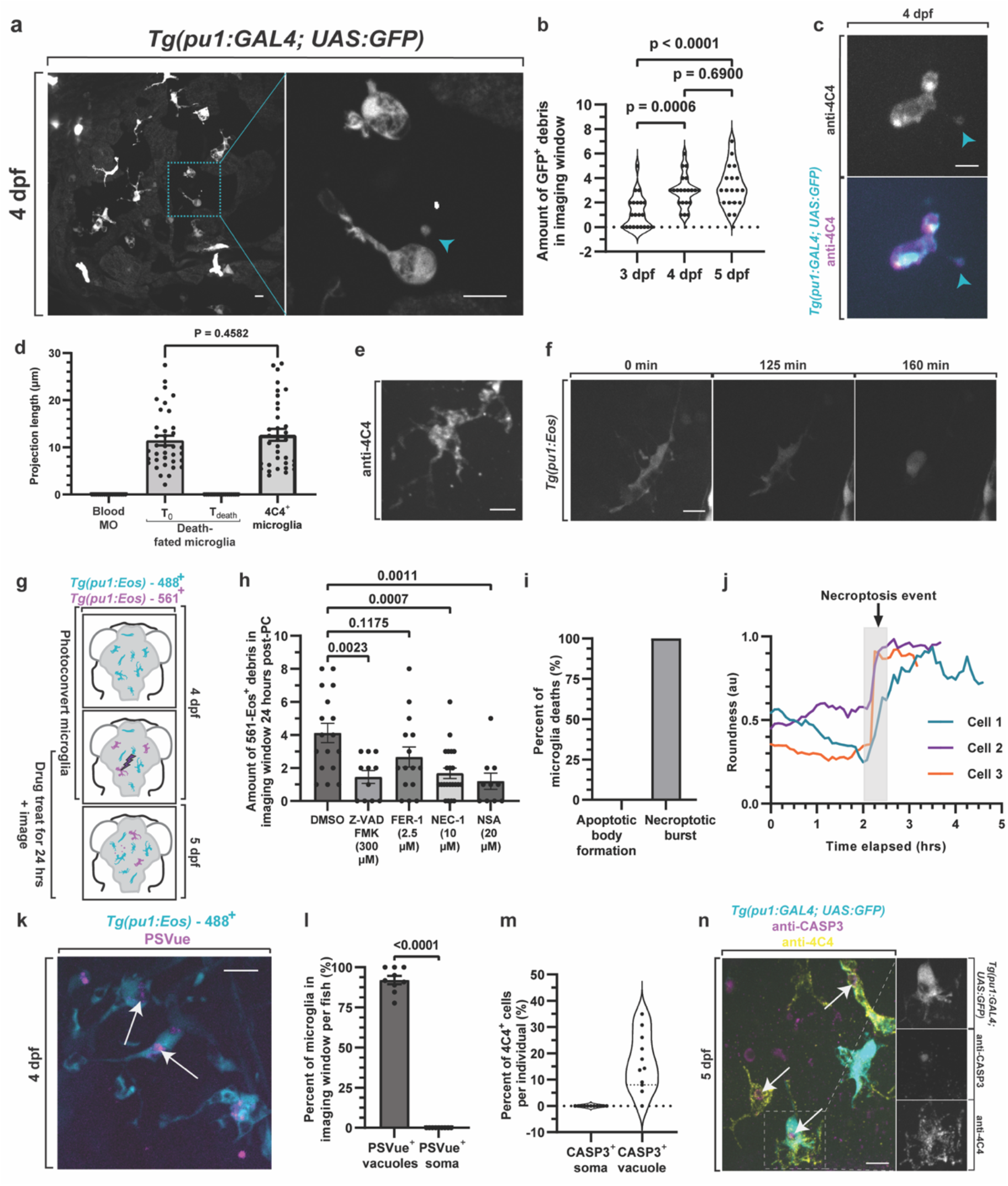
Microglia die via necroptosis in colonization stages. A-B. Confocal images of 4 dpf *Tg(pu1:GFP)* brains (A) and quantification (B) showing microglia debris (blue arrowhead) and intact microglia (p=0.1189 3 dpf vs 4 dpf, p=0.8368 4 dpf vs 5 dpf, p=0.0387 3 dpf vs 5 dpf, Post hoc Tukey test). C. Confocal images of 4 dpf *Tg(pu1:GFP)* brains labeled with anti-4C4 demonstrating GFP^+^; 4C4^+^ debris. D. Quantification of microglia projection length for blood macrophages (Blood MO), death-fated microglia at early timepoints and right before death from timelapse movies, and of microglia labeled with 4C4 at 4 dpf (p=0.4582 T_0_ death-fated microglia vs 4C4 microglia). E-F. Confocal image of 4C4 labeled microglia (E) and death-fated microglia (F) showing cellular morphology consistent with microglia identity at 4 dpf. G. Schematic of experimental paradigm (G) and quantification (H) of 561-Eos^+^ debris in the brain 24-hour after labeling showing Nec-1 and NSA reduce microglia debris (p=0.0023 DMSO vs Z-VAD FMK, p=0.1175 DMSO vs Fer-1, p=0.0007 DMSO vs Nec-1, p=0.0011 DMSO vs NSA, one-way ANOVA/Dunnett’s multiple-comparisons test). I. Quantification of the percentage of microglia deaths at XX dpf in 24h timelapse movies that cause apoptotic bodies vs necroptotic burst. J. Quantification of cell roundness in a 24 h movie from 4-5 dpf of microglia before it dies. K-L. Confocal images (K) and quantifications (L) of *Tg(pu1:Eos)* animals at 4 dpf that are treated with PsVue demonstrating PsVue labels vacuoles (arrow) but not cell bodies (p<0.0001, unpaired t-test). M-N. Quantifications (M) and confocal images of *Tg(pu1:GAL4;UAS:GFP)* animals labeled with Casp3 and 4C4 at 5 dpf (N) showing Casp3 is located in vacuoles (arrow) but not does not label the microglia cell bodies. Scale bar is 10 μm (a, c, e, f, k,n). One-way ANOVA (b, h), T-test (d,l). Additional supplemental material provided in Figure S1. Descriptive statistics represented in Table S1.

To investigate how this debris could occur, we generated movies of *Tg(pu1:GFP)* animals for 24 hours from 4-5 dpf. While shedding of GFP^+^ debris from living microglia is a possibility, we never saw such an event in 24-hour movies. Instead, we could detect that GFP^+^ microglia debris was caused by microglia cell death (Figure S1b, n=10 cells). Although our imaging used standard timelapse imaging approaches for zebrafish, we ruled out the possibility that the death was a result of imaging parameters by scoring the amount of death in animals that were timelapsed vs not timelapsed (Figure S1c,d). Staining for cleaved Caspase-3 further confirmed that the imaging itself was not causing death (Figure S1c). Consistent with other reports that imaged zebrafish microglia, imaging also did not cause an increase in microglia death or alter microglia abundance (Figure S1e,f,q,r). Further, the above data represents data from single-timepoints and timelapses, supporting the concept that the microglia death is occurring independent of imaging parameters. It is also possible that the GFP^+^ debris that is present were small cellular extensions from attached microglia but are undetectable with the 4C4 antibody or the cytosolic GFP in *Tg(pu1:GFP)*, but timelapse movies at 4 dpf demonstrated stationary GFP^+^ debris puncta that clearly separated from migratory GFP^+^ cells (Figure S1g,h, Movie S1). These timelapse imaging experiments revealed that GFP^+^ debris is quickly removed from the parenchyma (Figure S1i, Movie S1), thereby consistent with the idea that our quantifications of GFP^+^ debris is underreporting the total amount of microglia death.

Most developmental death is mediated by apoptosis^1^. To identify the molecular mechanism of this microglia death, we first treated animals with Z-VAD FMK from 4-5 dpf, a pan caspase inhibitor that is highly effective at blocking apoptosis in multiple cell types^22^. The Z-VAD FMK treatment was confirmed to reduce apoptosis at 5 dpf by scoring cCaspase3 in DMSO vs Z-VAD FMK (Figure S1j) To normalize the amount of microglia debris per cell, we next designed a strategy to label a specific number of microglia. To do this, we generated a *Tg(pu1:Eos)* animal, which expresses the photoconvertible protein, Eos, in microglia (Figure 1g)^23^. In this paradigm, we exposed three microglia to 405 nm light at 4 dpf and thereby photoconverted the Eos protein from green (488-Eos) to red (561-Eos) emission (Figure S1k, Movie S2). Scoring debris in DMSO vs Z-VAD FMK-treated animals at 5 dpf was statistically different (Figure 1h)(p=0.0023 DMSO (n=17) vs Z-VAD-FMK(n=11), Dunnett’s multiple comparisons test), indicating that the microglia death was partially dependent on apoptosis. To explore other molecular mechanisms of microglia debris, we then similarly treated animals with inhibitors of ferroptosis (Fer-1) and necroptosis (Nec-1)^24–26^. Treatment with necroptotic inhibitors, Nec-1, also reduced the amount of microglia debris while ferroptosis inhibitors did not (Figure 1h)(p=0.0007 DMSO (n=17) vs Nec-1 (n=22), p=0.0011 DMSO (n=17) vs NSA (n=10), Dunnett’s multiple comparisons test). To confirm our treatment of Nec-1 inhibits its well-defined Ripk1 target we tested activation of signaling pathways downstream of Ripk1. We could not identify antibodies that label Ripk1 and MLKL specifically in zebrafish tissue. We therefore tested if Nec-1 treatment perturbed other defined Ripk1-mediated signaling by determining if NFkB activation was disrupted^27^. We treated *Tg(nfkb:eGFP)* which uses NFkB activation domains to drive eGFP expression and detected reduced levels of eGFP in Nec-1 vs DMSO animals at 4 dpf (Figure S1l,m). This reduction is specific to Ripk1 inhibition because treatment with NSA did not change *nfkb:eGFP* (Figure S1l,m). To provide a complementary approach to test the role of necroptosis, we repeated the photoconversion paradigm with NSA, an inhibitor of MLKL that is downstream of Ripk1 and specific to necroptosis. Similar to Nec-1 treatment, NSA treatment also reduced the number of debris, consistent with the idea that embryonic microglia death is dependent on necroptosis (Figure 1h). While the amount of debris decreased after Nec-1 and NSA treatment, the overall number of microglia (as labeled by 4C4) after 24 hours of treatment with DMSO, Nec-1 or NSA treatments were indistinguishable (Figure S1n).

To further understand if embryonic microglia death was necroptotic or apoptotic and not just dependent on other necroptotic or apoptotic cells, we visualized microglia as they undergo death with *Tg(pu1:GFP)* animals in 24 hour time-lapse movies from 4-5 dpf. We scored the dynamic features of apoptotic cellular bodies that would be consistent with apoptosis^13,15,28^. In contrast, a rounding up of a cell followed by sudden disappearance of cytoplasmic labeling would be indicative of necroptosis^29^. Tracing of individual microglia every 5 minutes for 24 hours revealed that microglia death did not exhibit apoptotic cellular bodies (Figure 1i, N=5 animals, n=10 cells). Instead, most microglia that died halted their migration, retracted all of their cellular processes to reduce their area, rounded up and then shortly later, burst (Figure 1i,h, S1p, Movie S3). We further identified that microglia membranes at 4 dpf does not label with the phosphatidylserine reporter PSVue (Figure 1k,l, N=9 animals, n=92 cells), which is normally localized on apoptotic cells^30,31^. We complemented this approach with staining of cleaved-Caspase3, which is present in apoptotic cells. While cCaspase3 labeled embryonic brain cells at 4 dpf, we did not detect cCaspase3+ microglia (Figure 1m,n). Instead, cCaspase3 was only present within microglia phagosomes, similar to PSVue (Figure 1m,n). Together with the pharmacological inhibition, this strongly supports the idea that developmental microglia die via necroptosis. These experiments do not rule out that microglia could die via other cell death processes, but for this report, we focused on the necroptotic death of microglia.

To next investigate cell biological features that promote this microglia necroptosis, we tracked the lifespan of individual microglia until they died. To do this we utilized time-lapse imaging of the zebrafish brain to track individual microglia. In 4 dpf *Tg(pu1:GFP)* animals, we could detect microglia with cytosolic GFP labeling that contained GFP^-^ vacuole-like structures (Figure 2a). Transgenic animals for specific cell-types demonstrated the content of these vacuoles were astroglial (from *Tg(gfap:NTR-mCherry)*)^32^, neuronal (from *Tg(nbt:dsRed)*)^15^ and to a lesser extent oligodendroglial (from *Tg(sox10:mRFP)*)^33^ debris (Figure 2b, S2a,b). For this manuscript we define vacuoles as a descriptive term for XFP-negative inclusions within microglia. By collecting images that spanned the brain, we identified that vacuole containing microglia represented the majority of microglia in the developing brain (Figure S2c, N=5 animals). We fate mapped these microglia by imaging them in the brain every 5 minutes for 24 hours and then tracked microglia and the vacuoles in them at every time point from 4-5 dpf. Plotting each individual microglia in 24-hour periods demonstrated that microglia transitioned from non-vacuole containing to vacuole containing (Figure 2c,d). However, we could not detect microglia that transitioned from vacuole containing to non-vacuole containing over the 24-hour imaging period (Figure 2c,e, N=4 animals, n=12 cells). In these movies, we instead identified that all the microglia that died contained vacuole-like structures (Figure 2c,f, N=4 animals, n=10 cells), introducing the possibility that most embryonic microglia die before they completely digest efferocytic debris. To explore the possibility that overload of cellular debris caused microglia death, we measured the amount of vacuoles and size of vacuoles in movies of microglia 5 timepoints before they died and compared that to microglia that did not die in our imaging window. However, we did not detect any amount of load that predicted death between death-fated microglia and living microglia (Figure S2d,e).

**Figure 2.**
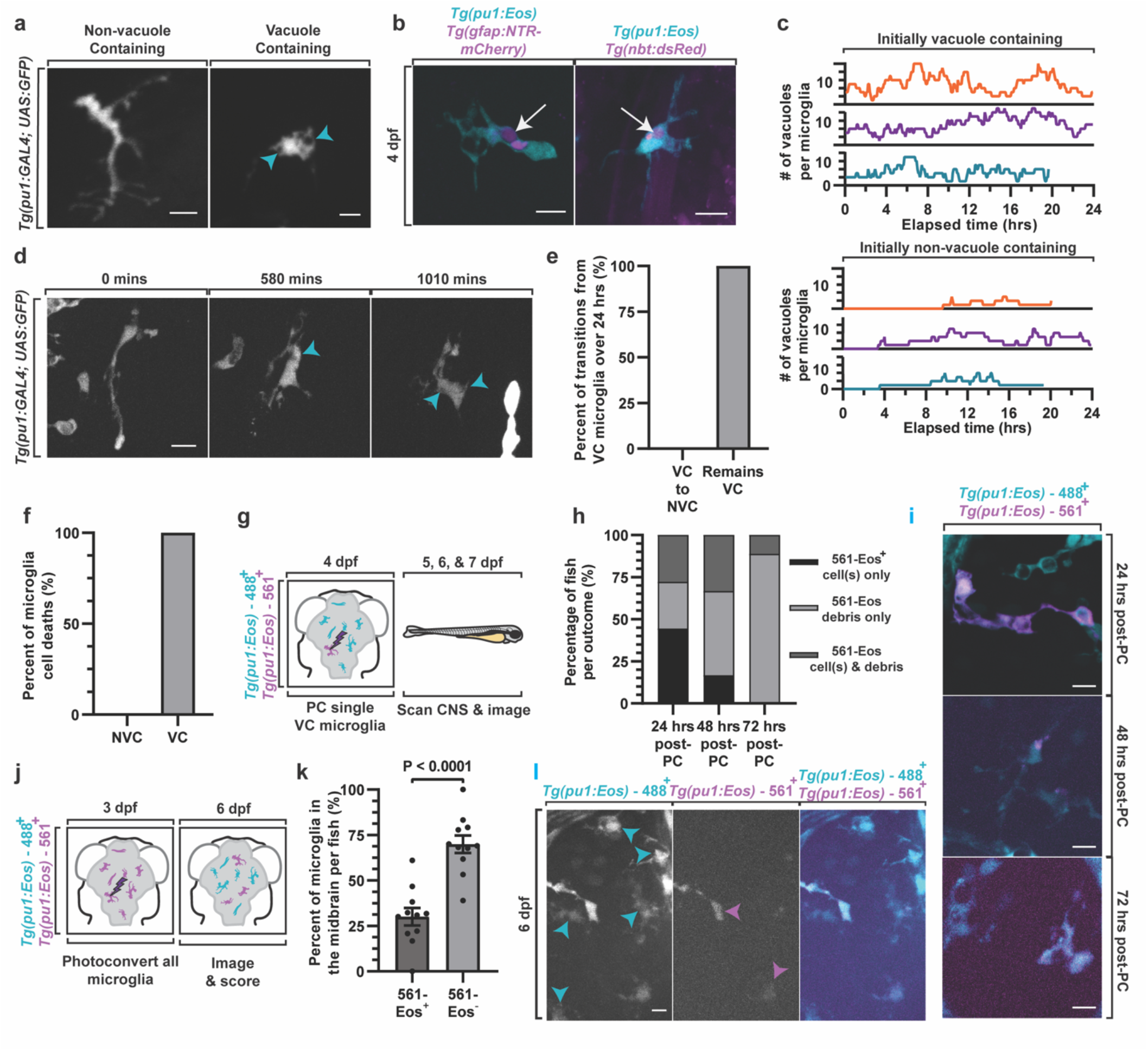
Developmental microglia have short lifespans after becoming vacuole-containing. A. Confocal images from a 24h timelapse movie from 4-5 dpf showing microglia in *Tg(pu1:gfp)* animals that contain vacuoles and microglia that do not have vacuoles. Arrowhead denotes vacuoles. B. Confocal images of *Tg(pu1:Eos); Tg(gfap:NTR-mCherry)* and *Tg(pu1:Eos); Tg(nbt:DsRed)* animals at 4 dpf demonstrating microglia that contain *gfap*^*+*^ and *nbt*^*+*^ debris. Arrow denotes debris within vacuoles. C-D. Quantifications (C) and confocal images (D) of vacuoles inside individual microglia from 24h timelapse from 4-5 dpf of *Tg(pu1:GFP)* animals. The number of vacuoles per microglia are scored every 5 minutes for 24 hours. Arrowhead denotes vacuoles. E-F. The percentage of total microglia that transition between different vacuole-containing states (E) and the state of cells before their death (F) in 24h timelapse movies. G-H. Schematic of experimental design (G), and quantifications (H) to test the fate of microglia. Quantifications in H demonstrate the percentage of occurrences that 561-Eos^+^ microglia in *Tg(pu1:Eos)* results in 561-Eos^+^ intact microglia, 561-Eos^+^ microglia debris or 561-Eos^+^ intact microglia and microglial debris 24, 48 and 72 hrs after labeling. I. Confocal images demonstrating outcomes that were quantified in H. J. Experimental paradigm schematic of experimental paradigm demonstrating that all microglia in the brain are labeled and then scored at 6 dpf. K. Quantification shows the percentage of microglia at 6 dpf that are labeled with 561-Eos and negative for 561-Eos (p<0.0001, unpaired t-test). L. Confocal images of microglia at 6 dpf in the experiment represented in J-K. Blue arrowheads represent microglia within Eos-561 labeling. Magenta arrowheads shows Eos-561^+^ microglia Scale bar is 10 μm (a,b,d,i,l). Additional supplemental material provided in Figure S2. Descriptive statistics represented in Table S1.

There seemed to be two simple hypotheses for these results: 1. either the completion of the digestion phase of microglia exceeds the 24-hour period or 2. developmental microglia die because they are mostly irreversibly vacuole containing. To explore these hypotheses, we tracked individual microglia over longer developmental periods. Using *Tg(pu1:Eos)* animals, we photoconverted a single microglia that contained vacuoles per animal at 4 dpf, then imaged single z-stacks at 1, 2, and 3 days after photoconversion in each animal (Figure 2g). After 24 hours, three outcomes occurred: 1. 27.78% of animals had only 561-Eos^+^ debris in the brain, 2. 27.78% of animals had 561-Eos^+^ debris and 561-Eos^+^ cells that contained vacuoles or 3. 44.44% of animals had 561-Eos^+^ cells that contained vacuoles, consistent with the idea that the microglia either died or divided (Figure 2h, N=18 animals). However, we could not detect 561-Eos^+^ microglia that did not contain vacuoles. 3 days after photoconversion we rarely detected intact 561-Eos^+^ microglia, but identified 561-Eos^+^ debris in the brain (Figure 2h, N=18 animals). While it is possible that detection of Eos was compromised by time or that the photoconversion paradigm promoted cell death, we confirmed with a second transgenic line, *Tg(sox10:Eos)*, that other long-lived, dividing cells^34,35^, like dorsal root ganglia cells, could still be visualized as intact 3 days after photoconversion (Figure S2f,g, N=8 animals, n=48 cells). These data are consistent with the likelihood that once developmental microglia have vacuoles, they die within 3 days.

Microglia are thought to be long-lived cells that initially populate the brain during development^3^. Our results, however, indicate that once embryonic microglia become vacuole-containing in the zebrafish brain, they have short lifespans (Figure 2h, N=18 animals). Scoring of microglia from 3-5 dpf also showed 76.65% ± 1.95% of microglia are vacuole-containing on a given day (Figure S2c), and thus may be expected to die. Therefore, we next explored the scale of microglia death in the embryonic brain. To do this, we photoconverted all microglia in the brain of *Tg(pu1:Eos)* animals at 3 dpf, and then scored the ratio of microglia that were photoconverted at 5 dpf (Figure 2j). These results indicated that on average 30.07% ± 4.81% of *pu1:Eos*^*+*^ cells in the brain were photoconverted (Figure 2k, N=11 animals, n=240 cells, S2h,i), supporting the idea that during peak stages of neural debris, microglia have short lifespans less than 3 days. We also expected to see a subset of 488-Eos^+^ microglia because embryonic microglia are still generated from new invading cells at this age^13^. Quantification of the number of 488-Eos+ (561-Eos-) cells supports this idea (Figure 2k). Scoring of all microglia demonstrated that the overall number of microglia also increases (Figure S2i), confirming previous findings that microglia continue to infiltrate the brain during colonization^13^. To ensure that photoconversion of all brain microglia did not impact overall population size, we scored the number of microglia in the brains of non-photoconverted and photoconverted animals at 6 dpf (Figure S2j). We found no difference in the number of microglia between photoconverted and non-photoconverted animals (S2j)

These results indicated that microglia debris is present when microglia are not only expanding but also phagocytic ^3,12,13,15,36– 39^, at least of apoptotic debris. It is unclear how necroptotic debris would be removed in the embryo. Therefore, we next asked which cell-type was clearing microglia necroptotic debris. To do this, we searched the brain for microglia debris encased in the different phagocytic cells. We first assayed microglial debris in *Tg(pu1:RFP)*;*Tg(glast:GFPcaax-TA-nucRFP)*^40^ and *Tg(pu1:GFP);Tg(sox10:mRFP)* at 4 dpf, which represent astroglia and neural crest/oligodendrocyte lineage cells that have been characterized as phagocytic^19,41,42^. However, we could not detect microglia debris encased in either cell-type. We therefore devised a strategy to visualize microglia debris within living microglia by using our *Tg(pu1:Eos)-*photoconversion paradigm to label microglia before they died and were cleared at 4 dpf (Figure 1g). We then collected images of the brain 24 and 48 hours after photoconversion when 561-Eos^+^ microglia debris was present. In this paradigm, we detected 561-Eos^+^ microglia debris within 488-Eos^+^ microglia of all photoconverted animals (Figure 3a,b,c, N=12 animals), revealing the concept of microglia cannibalism (Figure 3b).

**Figure 3.**
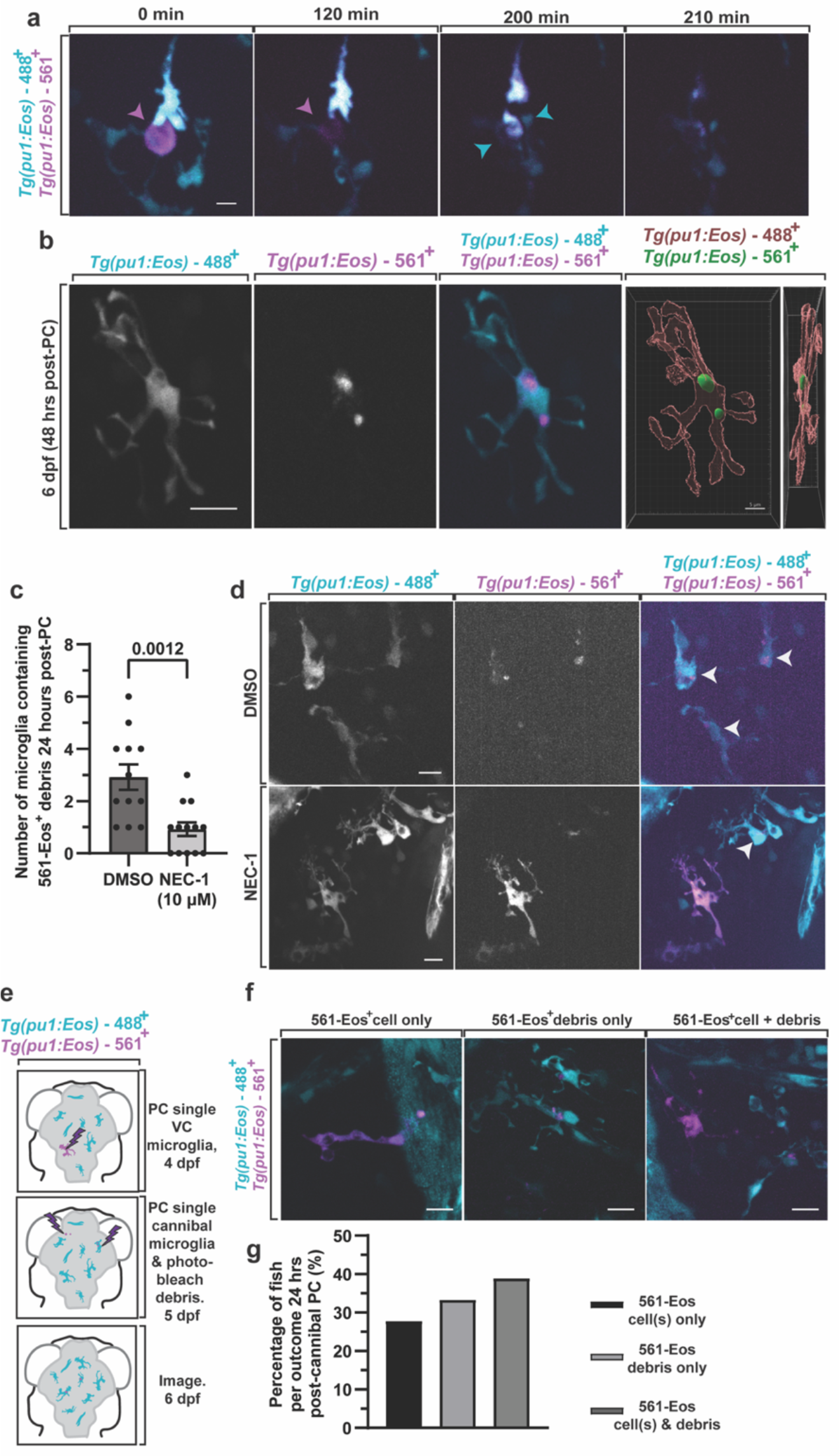
Microglia debris is cannibalized. A. Confocal images from 24h movie from 4-5 dpf of *Tg(pu1:Eos)* animals that had 561-Eos^+^ microglia that died and was cleared by 488-Eos^+^ microglia. Arrowheads denote a cell that undergoes death and is cannibalized. Arrowheads show microglia that dies and then is cleared by other microglia. B. Confocal images of *Tg(pu1:Eos)* at 24 hours form 4-5 dpf after labeling of 561-Eos^+^ microglia showing 561-Eos^+^ microglia debris within 488-Eos^+^ intact microglia. C-D. Quantification (C) and confocal images (D) of the amount of microglia that contained cannibalized 561-Eos^+^ debris in *Tg(pu1:Eos)* animals treated with DMSO and Nec-1 for 24 hours (p=0.0012 DMSO vs Nec-1, unpaired t-test). Arrowhead denotes cannibalistic microglia. E-F Schematic of experimental design (E), quantifications (G), and confocal images (F) to determine the fate of cannibalistic microglia. Quantifications score the fate (561-Eos^+^ intact microglia, 561-Eos^+^ microglia debris or 561-Eos^+^ intact microglia and microglial debris) of cannibalistic microglia. Scale bar is 10 μm (a,b,d,f). Additional supplemental material provided in Figure S3. Descriptive statistics represented in Table S1.

There were two likely hypotheses of how cannibalism occurred: 1. Microglia clear debris from dead microglia or 2. Microglia cannibalized portions of living microglia. To distinguish between these possibilities, we again labeled a random subpopulation of microglia with 561-Eos at 4 dpf and tracked the appearance of microglia debris within migrating microglia in 24h time-lapse movies. In these movies, the majority of cannibalism events showed a sudden disappearance of fluorescence in the dying microglia corresponding with the arrival of other microglia that instantly had 561-Eos^+^ puncta (Figure 3a). We did not detect microglia clearing of phagocytic cellular portions from other living microglia. To ensure this process was not a consequence of the photoconversion paradigm, we performed timelapse imaging of *Tg(pu1:GFP)* animals for 24 hours from 4-5 dpf. These movies revealed stationary and isolated GFP^+^ debris that rapidly disappeared when intact GFP^+^ microglia approached the debris (Figure 1f, S3a).

If microglia necroptosis was a causative event for microglia cannibalism, then inhibiting microglia death should reduce the amount of microglia cannibalism. We therefore scored the amount of cannibalism in DMSO vs Nec-1 treated animals at 5 dpf and detected that Nec-1 treated animals had less cannibalistic microglia than DMSO (Figure 3c,d)(p=0.0012 DMSO (N=12) vs NEC-1 (N=13), unpaired t-test), consistent with the hypothesis that microglia cannibalism is driven by microglia death. Taken together, this introduces a potential paradox that once developmental microglia become phagocytic, they die, which produces debris that is eventually cannibalized by other microglia – such cannibalistic microglia are then vacuole-containing and thus may be expected to die, ultimately driving turnover of embryonic microglia.

To explore this paradox, we tracked the fate of embryonic microglia that were cannibalistic. In this paradigm, we first photoconverted an individual microglia in *Tg(pu1:Eos)* animals at 4 dpf and then selected animals at 5 dpf that had 561-Eos^+^ debris inside 488-Eos^+^ microglia but did not contain intact 561-Eos^+^ cells. We then bleached all the 561-Eos^+^ debris in the brain of these animals, then photoconverted Eos to label the cannibalistic microglia with 561-Eos. 24 hours later, these animals were then scored for the presence of 561-Eos^+^ debris and/or cells in the brain (Figure 3e,f,g N=18 animals). Control animals that were not rephotoconverted demonstrated negligible levels of photoconverted-debris supporting the idea the 561-Eos was bleached in the experiment paradigm (Figure S1b,c). In animals where cells were photoconverted a second time, we identified 33.33% of cannibalistic microglia generated 561-Eos^+^ debris in the brain, 38.89% of cannibalistic microglia caused 561-Eos^+^ cells and 561-Eos^+^ debris and 27.78% of cannibalistic microglia produced 561-Eos^+^ cells (Figure 3g, N=18 animals), consistent with the hypothesis that most cannibalistic microglia die.

We sought to further explore the paradox that most phagocytic microglia die, leading to debris that must be cannibalized, which then produces more phagocytic microglia that are destined to die (Figure 4a). Two predictions could be made based on that hypothesis, 1. decreasing microglia phagocytosis should reduce microglia debris and cannibalism and 2. increasing neural debris and thereby microglia phagocytosis should increase microglia debris and cannibalism. We first reduced phagocytosis by treating *Tg(nbt:dsRed); Tg(pu1:GFP)* animals with L-SOP from 4-5 dpf, which inhibits phosphatidylserine-dependent phagocytosis that microglia utilize to clear dead neurons^28^. Animals treated with L-SOP from 4-5 dpf, had a reduced number of vacuole-containing microglia compared to DMSO-treated animals (Figure 4b,c)(DMSO (N=6 animals, n=132 cells), L-SOP (N=6 animals, n=171 cells, p=0.0034 DMSO vs L-SOP, unpaired t-test), confirming the treatment can inhibit phagocytosis^28^. L-SOP treated animals also had less GFP^+^ microglial debris than DMSO (Figure 4d,e)(p=0.0005, DMSO (N=6 animals, n=132 cells), L-SOP (N=6 animals, n=171 cells, unpaired t-test), indicating that inhibiting phagocytosis leads to less microglia death. To test if this also reduced microglia cannibalism, we photoconverted 3 microglia in *Tg(pu1:Eos)* at 4 dpf, and then scored 561-Eos+ debris in 488-Eos+ microglia. Animals were either treated with water or L-SOP from 4 dpf to 5 dpf. L-SOP treated animals had 1.125±0.295 cannibalistic microglia compared to 3.000±0.5345 water controls, suggesting that limiting phagocytosis reduces microglia cannibalism (p=0.0072, Water (N=7 animals) vs L-SOP (N=8 animals), unpaired t-test)(Figure 4f,g).

**Figure 4.**
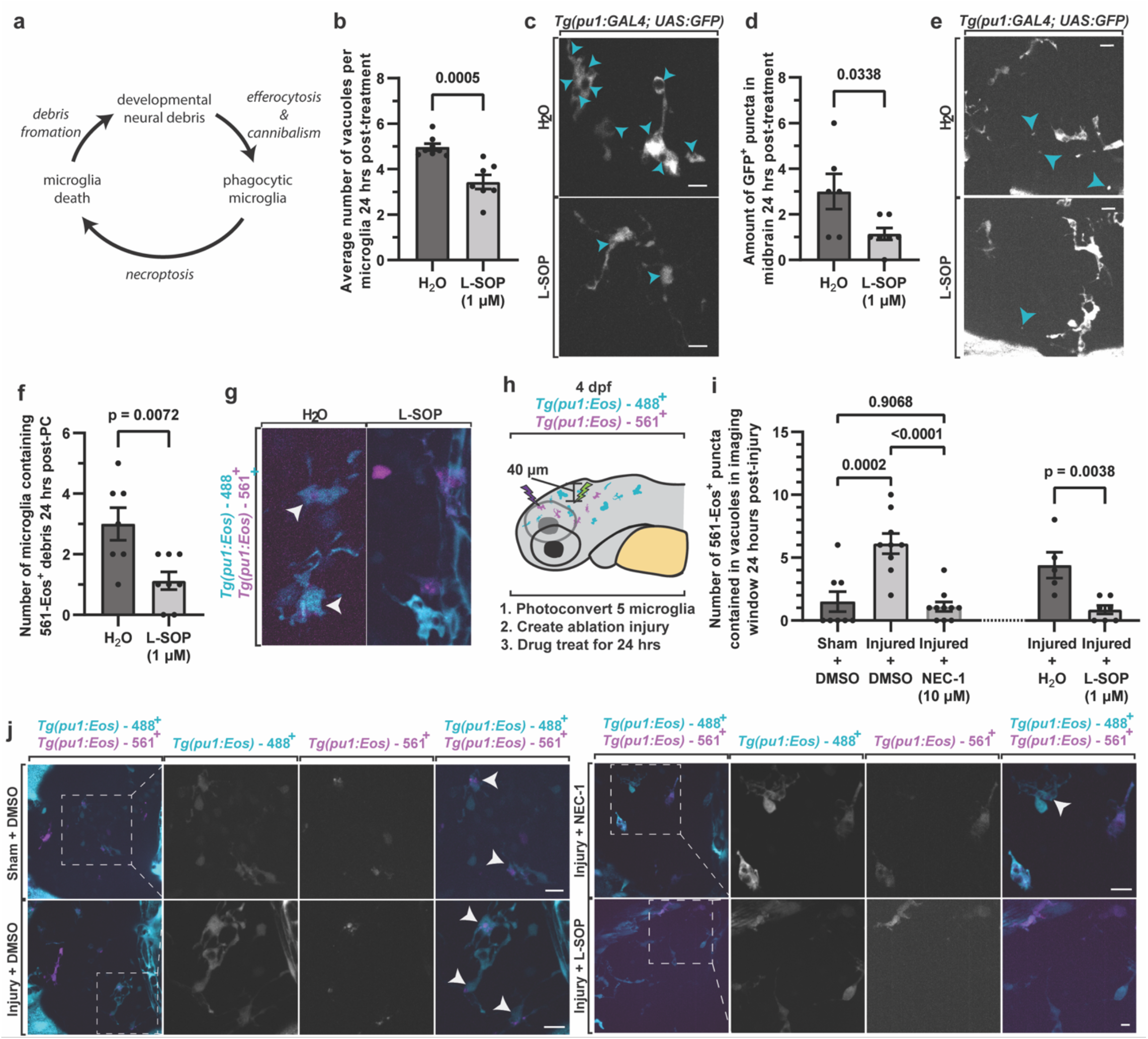
Phagocytosis and cannibalism shortens the lifespan of microglia. A. Schematic of the paradox that involves phagocytosis, necroptosis, and cannibalism. B. Quantifications of the average number of vacuoles in microglia at 5 dpf (p=0.0338 H_2_O vs L-SOP, unpaired t-test). C. Confocal images of *Tg(pu1:GFP)* animals at 5 dpf showing a reduction of vacuoles in microglia. Arrow denotes vacuoles). D. Quantification of the amount of microglia debris in *Tg(pu1:GFP)* animals treated with H_2_O (control) or L-SOP (p=0.0005, H_2_O vs L-SOP, unpaired t-test). E. Confocal images demonstrating a reduction in the amount of microglia debris from D. Arrowheads denote vacuoles. F. Quantifications of the amount of microglia that contain 561-Eos in water for L-SOP treatments. G. Confocal images of cannibalistic microglia at 5 dpf from treatments in F. H-I. Schematic of experimental design (H) and quantifications of cannibalistic microglia after laser-induced brain injury (I). Quantifications show the number of 561-Eos^+^ puncta in 488-Eos^+^ intact microglia 24 hours after sham injuries or laser-injuries in *Tg(pu1:Eos)* animals treated with DMSO vs Nec-1 and H_2_O vs L-SOP for 24 hours after the injury (p=0.0002 Sham+DMSO vs Injured+DMSO, p<0.0001 Injured+DMSO vs Injured+Nec-1, p=0.9068 sham+DMSO vs injured+Nec-1, Post hoc Tukey test). J. Confocal images from the experiment in H-I showing changes in microglia cannibalism at 5 dpf after injury in Nec-1 and L-SOP treatments. Arrowheads denote cannibalistic microglia. Scale bar is 10 μm (c,e,j). Additional supplemental material provided in Figure S4. Descriptive statistics represented in Table S1.

To test the paradox by increasing neural debris, we created injuries in the brain to non-microglia cells to generate more neural debris that is cleared by microglia. Injuries were created with a 532 nm single-pulsed laser^16,43,44^. As a sham-injury control, the 641 nm laser was exposed to a similar size injury site. We first established that our injury paradigm created neuronal debris in *Tg(nbt:dsRed); Tg(pu1:Eos)* by scoring the amount of *dsred*^*+*^ puncta in the brain after the injury at 5 dpf (Figure S4a). Despite an average of 4.667±12.69 neuronal *dsred*^*+*^ debris per imaging window, the injury paradigm did not cause detectable 488-Eos^+^ microglia debris immediately after the injury (Figure S4a,b)(N=6 animals, p<0.0001, dsRed^+^ debris vs. Eos^+^ debris, unpaired t-test). Consistent with the expansion of microglia after injury, 24 hours later we visualized an increase in microglia compared to sham-injured animals (Figure S4c)(Sham (N=5 animals) vs Injury (N=4 animals), p=0.0576, unpaired t-test). Having established the injury paradigm, we repeated it in *Tg(pu1:Eos)* animals that had 5 microglia in the brain photoconverted and thereby labeled with 561-Eos^+^ at 4 dpf (Figure 4h). After 24 hours, we quantified that the number of 561-Eos^+^ microglia debris present in the brain was in higher abundance in injured animals compared to sham-injured animals (Figure 4i,j)(Sham/DMSO (N=8) vs Injury/DMSO (N=9), p=0.0002, Post hoc Tukey test). The vast majority of this microglia debris was cannibalized by 488-Eos^+^ microglia.

If the increase in microglia phagocytosis of neuronal debris is causing microglia to die, and thereby causing more microglia cannibalism, then blocking microglia death or microglia phagocytosis should reduce the amount of microglia debris and microglia cannibalism. We, therefore, first tested if Nec-1 treatment reduced the number of cannibalistic microglia in the injury paradigm. In Nec-1 treated animals, the amount of 561-Eos^+^ microglia debris at 5 dpf returned to the level of sham-injured animals (Figure 4i,j)(Injury/Nec1 (N=10) vs Injury/DMSO (N=9), p<0.0001, Post hoc Tukey test). Microglia cannibalism also returned to the level of the uninjured animals, consistent with the hypothesis that phagocytic microglia death and cannibalism is impacted by necroptosis. One caution in the interpretation of the Nec-1 treatment results is that necroptotic microglia could be a direct consequence of the injury and not secondary to an increase in phagocytic microglia. To distinguish between such possibilities, we repeated the paradigm in animals treated with L-SOP from 4-5 dpf. In L-SOP treated animals that were injured, 0.8571±0.8997 561-Eos^+^ puncta were present in vacuoles at 5 dpf, compared to 4.400±1.030 in control animals (p=0.0038, unpaired t-test H_2_O (N=5 animals) vs L-SOP (N=7 animals) (Figure 4i,j). The simplest explanation for this collective data is that efferocytosis by microglia results in a shorted lifespan via necroptosis.

Together, our data shows that microglia that contain phagocytic debris in vacuoles have short lifespans that rarely exceed 3 days. Upon their death, developmental microglia debris is cannibalized by other microglia. The phagocytic nature of cannibalistic microglia then leads to their death and more debris that must be phagocytosed. This paradoxical process thereby establishes a cycle. In development, the paradox could be started by the apoptotic death of neural cells, which are cleared by microglia. This hypothesis may explain why inhibition of apoptosis and necroptosis reduces microglia death even though we could not detect apoptotic-based death in our imaging. This work, with others, also establishes that microglia debris is cleared by astrocytes and microglia in specific contexts^19^. Beyond the universal principle that cells overexpand during development and then die via apoptosis^1,2^, this work reveals an additional mechanism that produces developmental debris, induced by sacrificial microglia that function to clear apoptotic debris. This work further introduces an additional attribute of microglia expansion in the embryo. In addition to previous studies that demonstrated microglia expansion is balanced by infiltration of progenitors and cell division of existing microglia^13^, the death of microglia could also contribute to the overall abundance of microglia in the embryo. Future studies that investigate how these different attributes of the embryonic microglia pool are connected will be important. Given the data here and evidence of necroptotic microglia in disease contexts^17,18^, it will be intriguing to investigate if microglia cannibalism and a subsequent paradox that is driven by microglia cell death, is a hallmark of neuropathologies through ages.

### Limitations to Results

This work supports the idea that microglia die via necroptosis but does not rule out the possibility that additional modes of cell death contribute to microglia death. We use the term vacuoles above to describe a feature in microglia. These vacuoles contain debris from different neural cells and thus likely represent phagosomes. However, without testing specific phagosome markers to confirm that and to avoid giving them a biological function that was not tested (i.e. phagosome, lysosome), we elected to describe them with a more descriptive term, vacuoles. Future work will need to investigate the deeper molecular mechanism and biological significance of microglia cannibalism.

## AUTHOR CONTRIBUTIONS

HG performed the analysis, experimentation, writing, and editing of the manuscript. ZS provided reagents and advice on cell death experiments. CJS conceived, wrote and edited the manuscript and supervised and funded the project.

## ACKNOWLEDGEMENTS

We thank Beth Stevens, Chris Bennett, Mariko Bennett and Siyuan Zhang for helpful feedback and current and previous members of the Smith Lab for insightful discussions. We especially thank Abigail Zellmer for her efforts in constructing *Tg(pu1:Eos)nt200*. We also thank 3i for imaging related questions, Sara Cole in the NDiiF Optical Microscopy Core for help with IMARIS analysis, and Deborah Bang, and Matthew Lewis for zebrafish housing and upkeep.

## MATERIALS AND METHODS

### Contact for Reagent and Resource Sharing

All data collected for the study are included in the figures and supplemental data. Reagents are available from the authors upon request.

### Experimental Model and Subject Details

#### Ethics Statement

Experimental procedures followed the NIH guide for the care and use of laboratory animals. The University of Notre Dame IACUC which is guided by the United States Department of Agriculture, the Animal Welfare Fact (USA) and the Assessment and Accreditation of Laboratory Animal Care International approved all animal studies under protocol 22-07-7322 to Cody J. Smith.

#### Contact for Reagent and Resource Sharing

All data collected for the study are included in the figures and supplemental data. Reagents are available from the authors upon request.

### Experimental Model and Subject Details

Zebrafish strains in this study include: AB, *Tg(pu1:GAL4, UAS:GFP)*^*zf149*,15^, *Tg(gfap:NTR-mCherry*)^sc059,45,46^, *Tg(nbt:dsRed)*^*zf148*,15^, *Tg(pu1:Eos)*^*nt200*,23^, *Tg(sox10:Eos)*^*w9*,34^, *Tg(sox10:mRFP)*^*vu234*,33^, *Tg(slc1a3b:myrGFP-P2A-H2AmCherry*)^40^, *Tg(pu1:GAL4, UAS:RFP)*^*hdb2*,15^, *Tg(nfkb:eGFP)*^47^. Only germline transgenic lines were used in this study. To produce embryos, pairwise matings were used. Animals were raised at 28°C in egg water in constant darkness and staged by hours or days post fertilization (hpf and dpf), confirmed by observation of developmental milestones^48^. Embryos were used for all experiments.

## Method Details

### *In vivo* imaging

Animals were anesthetized using 3-aminobenzoic acid ester (Tricaine), covered in 0.8% low-melting point agarose, and mounted dorsally in glass-bottomed 35 mm petri dishes^49^. A spinning disk confocal microscopes custom built by 3i technology (Denver, CO) that contains: Zeiss Axio Observer Z1 Advanced Mariana Microscope, X-cite 120LED White Light LED System, filter cubes for GFP and mRFP, a motorized X,Y stage, piezo Z stage, 20X Air (0.50 NA), 63X (1.15NA), 40X (1.1NA) objectives, CSU-W1 T2 Spinning Disk Confocal Head (50 μm) with 1X camera adapter, and an iXon3 1Kx1K EMCCD or Teledyne Prime 95B cMOS camera, dichroic mirrors for 446, 515, 561, 405, 488, 561,640 excitation, laser stack with 405 nm, 445 nm, 488 nm, 561 nm and 637 nm with laser stack FiberSwitcher, photomanipulation from vector high speed point scanner ablations at diffraction limited capacity, Ablate Photoablation System (532 nm pulsed laser, pulse energy 60J @ 200 HZ) was used to acquire images^49^. Images in time-lapse microscopy were collected every 5 min for 24 hours. Images were processed with Adobe Illustrator, ImageJ, and IMARIS. Only brightness and contrast were adjusted and enhanced for images represented in this study.

### Immunohistochemistry

The primary antibody used in the confirmation of microglia debris in the brain was anti-cleaved Caspase3 (1:700, BD Biosciences) and anti-4C4 (1:50, mouse, Seiger, Becker and Becker Laboratories)^20^. The secondary antibody used was Alexa Fluor 647 goat anti-mouse (1:600, Thermo Fisher, A-21235). Staining was performed using the protocol by Nichols and Smith^49^. Larvae were fixed at 3 dpf, 4 dpf, and 5 dpf in fresh 4% paraformaldehyde in 0.1% PBS Triton-X.

### Photoconversion experiments

#### Single-cell photoconversions

*Tg(pu1:Eos)* embryos were grown to 4 dpf. Pre-conversion confocal *z*-stack images were taken in the 488 nm/GFP filter set and 561 nm/RFP filter set to confirm there was no nonspecific photoconversion. Single, vacuole-containing *pu1*^*+*^ cells were then photoconverted using a 5-ms 405-nm-laser pulse guide to the midbrain with mVector (3i) in the midbrain at 4 dpf following the single-cell photoconversion protocol described in Green and Smith^50^. Following photoconversion, confocal *z*-stack images were taken in the 488 nm/GFP filter set and 561 nm/RFP filter set to confirm expression of the converted Eos^+^ protein. Photoconverted animals were then grown to and imaged at 5 dpf, 6 dpf, and 7 dpf. Confocal 140 um *z*-stack images were taken of the dorsally mounted midbrains and the trunks of the fish were manually scanned for presence of 561-Eos^+^ cells and debris. We categorized the fish based on the presence of 561-Eos^+^ debris and 561-Eos^+^ cells.

#### Cannibal tracking

*Tg(pu1:Eos)* embryos were grown to 3 dpf. Pre-conversion confocal *z*-stack images were taken in the 488 nm/GFP filter set and 561 nm/RFP filter set to confirm there was no nonspecific photoconversion. Single, vacuole containing *pu1*^*+*^ cells were then photoconverted using a 5-ms 405-nm-laser pulse in the midbrain at 3 dpf following the single-cell photoconversion protocol above. Following photoconversion, post-conversion confocal *z*-stack images were taken in the 488 nm/GFP filter set and 561 nm/RFP filter set to confirm expression of the converted Eos^+^ protein. Photoconverted animals were then grown to and imaged at 4 dpf. Confocal 140 um *z*-stack images were taken of the dorsally mounted midbrains and the trunks of the fish were manually scanned for presence of 561-Eos^+^ cells and debris. Fish with 561-Eos^+^ cells at 4 dpf were removed from the experiment. Cannibal microglia were identified as 488-Eos^+^ microglia containing 561-Eos^+^ debris in the remaining fish. One cannibal microglia was then selected per fish and photoconverted. All other 561-Eos^+^ debris present was photobleached using as many 5-ms 561-nm-laser pulses as necessary. Fish were grown to and imaged at 5 dpf, 6 dpf, and 7 dpf. We categorized the fish based on the presence of 561-Eos^+^ debris and 561-Eos^+^ cells. To confirm the bleaching distinguished the 561^+^-Eos signal, a subset of animals was photobleached but not re-photoconverted.

### Injury

*Tg(nbt*:*dsRed);Tg(pu1*:*Eos)* and *Tg(pu1:Eos)* 4 dpf animals were anesthetized using 0.02% 3-aminobenzoic acid ester (Tricaine) in egg water. Fish were then dorsally mounted in 0.8% low-melting point agarose solution, arranged laterally on a 10 mm glass-coverslip–bottom Petri dish, and placed on the microscope anterior to posterior. Injured were performed in the midbrain. Specific site of laser-induced injury was determined by bringing the skin of the fish above the brain into focus and using the piezo Z stage to move 40 μm below the surface of the skin. This area was marked and brought into a focused ablation window. Upon focusing the targeted region, we double-clicked on a *dsRed*^*+*^ region using a 4 μm cursor tool. All laser parameters used are specific to our confocal microscope. Specific parameters include Laser Power (1), Raster Block Size (1), Double-Click Rectangle Size (4), and Double-Click Repetitions (4). After injury, fish were released from the agarose and treated with 1% DMSO or Nec-1. Sham-injured fish were followed the same procedure but were expose to single pulses of 561 nm laser with mvector rather than the Ablate! Laser.

### Chemical treatments

#### Cell death inhibitors

The chemical reagents used were Z-VAD-FMK^22^, Ferrostatin-1 (Fer-1; Sigma, SML0583)^26^, Necrostatin-1 (Nec-1; MedChemExpress LLC, HY-15760)^25^, Necrosulfamide (NSA; Tocris Bioscience)^24^. Stock solutions of 20 mM Z-VAD-FMK, 2.5 mM Fer-1, 10 mM Nec-1, and 10 mM NSA were stored at −80°C dissolved in DMSO. Working solutions were diluted with PTU to 300 μM for Z-VAD-FMK treatments, 2.5 μM for Fer-1 treatments, 10 μM for Nec-1 treatments^51^, and 20 μM for NSA treatments. All embryos were incubated in egg water until 24 hpf and incubated with PTU until desired treatment time. Fish were treated at 4 dpf, immediately after photoconversions. Control fish were incubated with 1% DMSO in PTU.

#### L-SOP treatment

The chemical reagent used for this study was O-Phospho-L-serine (L-SOP; Sigma, P0878-10MG). Stock solutions were dissolved in H_2_O to a concentration of 1 mM. Working solutions were diluted to 1 μM with PTU^28^. All embryos were incubated in egg water until 24 hpf and incubated with PTU until desired treatment time. Fish were bathed at 4 dpf with 1 μM L-SOP dissolved in egg water for 24 hours. Control fish were incubated with H_2_O.

#### PSVue Labeling

For imaging determining if microglia express phosphatidyl serine, embryos were bathed in a 1:250 solution of PSVue^®^ 643 (PSVue; Polysciences, Inc.) diluted in egg water for 1 hour prior to imaging.

### Quantification and statistics

3i Slidebook software was used to generate composite z-stack images of microglia. All individual z-stack images were sequentially observed. IMARIS (Notre Dame Imaging Core) was used to create 3D surface renderings of microglia. All graphical data represent both the mean and individual values used in each experiment unless otherwise noted. All quantifications were performed using various plug-ins available in FIJI (ImageJ) and Microsoft Excel. GraphPad Prism (version 8) software was used to perform all statistical analysis.

No statistical methods were used to predetermine sample sizes, however all sample sizes are informed by previous publications.

All statistical tests were run with biological replicates, not technical replicates. Healthy animals were randomly selected for experiments. No data points were excluded from analysis. Data distribution was assumed to be normal, but this was not formally tested. Unless otherwise indicated, data collection and analysis were performed blind to the conditions of the experiments. Each experiment was repeated at least twice with similar results.

#### Quantification of debris

GFP^+^ debris, 488-Eos^+^ debris, 561-Eos^+^ debris, *gfap*^+^ debris, *nbt*^+^ debris, and *sox10*^+^ debris were counted manually in Slidebook and ImageJ across all consecutive images in *z*-stacks of the midbrain. 488-Eos^+^ microglia were considered cannibalistic if they contained 561-Eos^+^ debris.

#### Quantification of vacuoles

24 hour timelapse movies were taken of 4 dpf *Tg(pu1:GAL4; UAS:GFP)* fish. Individual microglia were manually tracked at every 5 minute timepoint of the timelapses. At each timepoint, the number of vacuoles an individual microglia contained was counted by going through images in *z*-stacks and was supported by the use of Slidebook’s 4D volume view feature. Vacuoles were counted at every timepoint until the microglia left the capture window. Vacuoles were defined as GFP^-^ inclusions of any size that were completely contained within GFP^+^ microglia.

#### Cell morphology quantifications

Analysis of microglia area and roundness were used to describe changes in microglia morphology. Microglia area and roundness were measured at every 5-minute time point before death using the trace feature on ImageJ. ImageJ calculates roundness using the formula:

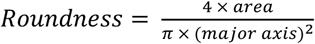

#### IMARIS

3D surface reconstructions were generated using IMARIS. The surface tool was use to generate surface renderings from confocal stacks taken with 1 um step sizes. Only brightness and contrast were adjusted.

### Softwares

ImageJ and Slidebook were used to produce and process confocal images. Graphpad prism was used to generate all graphs and statistical analysis. Adobe Illustrator was used to compile the figures and p1.

## DATA AVAILABILITY

All data collected for this study are included in the figures and supplementary material.

## SUPPLEMENTAL FIGURES

**Figure S1.**
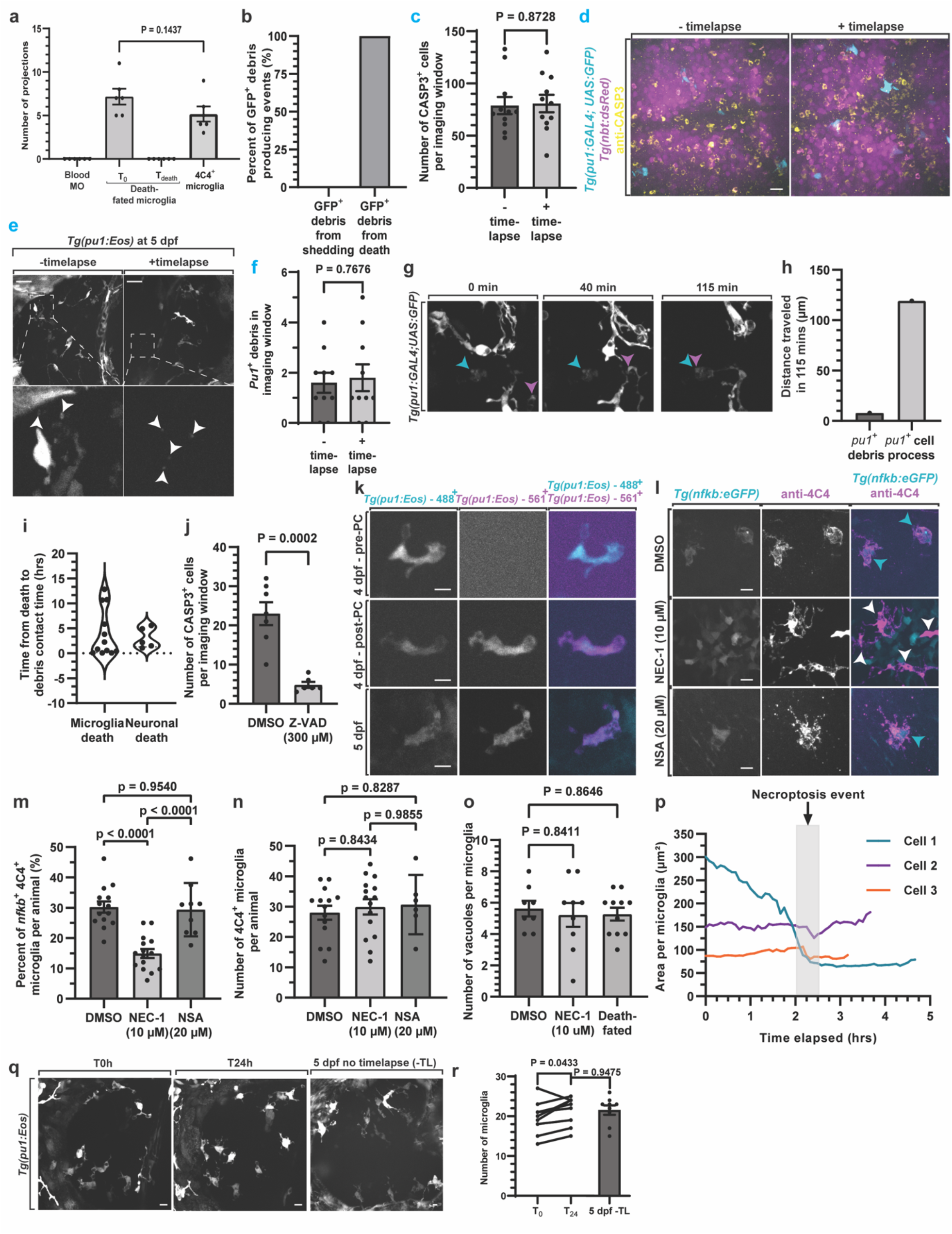
Supplemental material for Figure 1. A. Quantification of microglia projection abundance for blood macrophages (Blood MO), death-fated microglia at early timepoints and right before death from timelapse movies from 4-5 dpf, and of microglia labeled with 4C4 at 4 dpf (p=0.1437 T0 death-fate microglia vs 4C4 microglia). B. Quantification of the amount of debris causing events that result from shedding vs cell death in 24-hour timelapse movies of 4 dpf *Tg(pu1:GFP)* animals. C. Quantification of CASP3+ cells in animals that were time-lapses for 24 hours vs. not timelapsed from 4-5 dpf. (p=0.8728 timelapsed vs not timelapsed). G. Confocal images from a 24-hour timelapse from 4-5 dpf of *Tg(pu1:GFP)* animals demonstrating debris that is distinct from intact microglia (blue arrowhead) that disappears after intact microglia (magenta arrowhead) migrates across it. H. Quantifications from D demonstrating that *pu1*^*+*^ debris is not migratory like *pu1*^*+*^ cells and thereby not physically connected. I. Quantification from a timelapse from 4-5 dpf of *Tg(pu1:GFP)* (microglia death) and *Tg(nbt:dsRed)* (neuronal death) animals showing how quickly debris is cleared in the brain. J. Quantification of the number of CASP3+ cells in animals treated with DMSO vs Z-VAD (p=0.0002). K. Images from a confocal microscope of 4 dpf *Tg(pu1:Eos)* animals before and after exposure to 405 nm. Note that absence of *Tg(pu1:Eos)* -561^*+*^ before photoconversion. L. Confocal images of *Tg(nfkb:GFP)* animals at 5 dpf stained with 4C4 after treatment for 24 hours of DMSO, NEC-1, and NSA. M. Quantification of the percent of *nfkb*+; 4C4+ microglia in *Tg(nfkb:GFP)* animals at 5 dpf stained with 4C4 after treatment for 24 hours of DMSO, NEC-1, and NSA. (p<0.0001 DMSO vs NEC-1, p=0.9540 DMSO vs NSA).. N. Quantification of the abundance of microglia (N) and number of vacuoles in those microglia (O) of 5 dpf animals treated for 24 hours with DMSO, NEC-1, and NSA. P. Quantification of the area of microglia during cell death event from 4-5 dpf. Q-R. Confocal images of *Tg(pu1:GFP)* animals (Q) and quantifications of microglia abundance (R) at timepoint 0hr and 24hr from a 24-hour timelapse and non-timelapsed animals demonstrating that the number of microglia is not impacted by 24-hour timelapses. Note that cells rapidly decrease area just before the necroptosis event. Scale bar is 25 μm (d,e). Scale bar is 10 μm (k,l). Descriptive statistics represented in Table S1.

**Figure S2.**
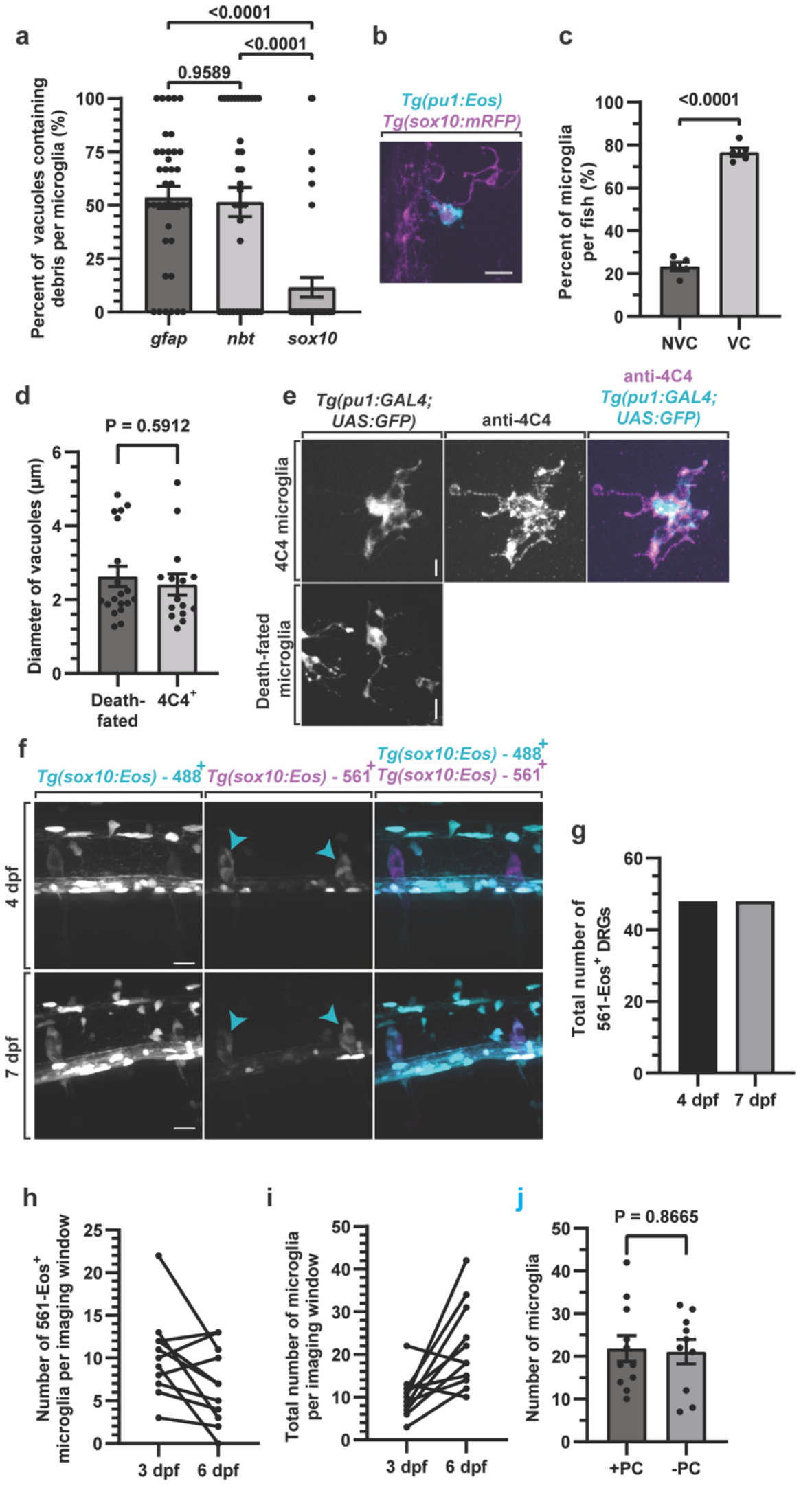
Supplemental material for Figure 2. A. Quantification of the percentage of vacuoles in microglia that enclose each type debris from *Tg(gfap:NTR-mCherry), Tg(nbt:dsRed)* and *Tg(sox10:mRFP)* animals at 4 dpf (p=0.9589 *gfap* vs *nbt*, p<0.0001 *gfap* vs *sox10*, p<0.0001 *nbt* vs *sox10*, Post hoc Tukey test). B. Confocal images of *Tg(pu1:Eos); Tg(sox10:mRFP)* animals at 4 dpf showing *sox10*^*+*^ debris within *pu1*^*+*^ microglia. C. Quantification of the percentage of microglia per 4 dpf animals that are non-vacuole (NVC) vs vacuole containing (VC)(p<0.0001 NVC vs VC, Fisher’s exact test). D. Quantification of the diameter of microglia vacuoles from 24 hour timelapse movies of *Tg(pu1:GFP)* animals at 4 dpf (death-fated). Such microglia were compared to microglia that were labeled with 4C4 at the corresponding age (p=0.5912, t-test). E. Confocal images of death-fated microglia and microglia stained with 4C4 that were used to generate D. F. Confocal images of *Tg(sox10:Eos)* animals that were photoconverted at 4 dpf and imaged at 4 and 7 days, demonstrating the Eos photoconversion is stably detected at least 3 days after photoconversion. G. Quantifications from F that demonstrate that photoconversion causes stable labeling of Eos^+^ cells. H. Quantification of the abundance of 561-Eos^+^ microglia in *Tg(pu1:Eos)* animals that had all microglia photoconverted a 3 dpf and then quantified at 6 dpf. Each data point with connect line represent a single animal. Note the decrease in 561-Eos+ microglia. I. Quantification of animals depicted in H and total number of microglia is quantified. Note that the overall abundance of microglia increases. J. Quantification of number of microglia in photoconverted (+PC) and non-photconverted (-PC) animals at 6 dpf (p=0.8665, t-test). Scale bar is 10 μm (b,e,f). Descriptive statistics represented in Table S1.

**Figure S3.**
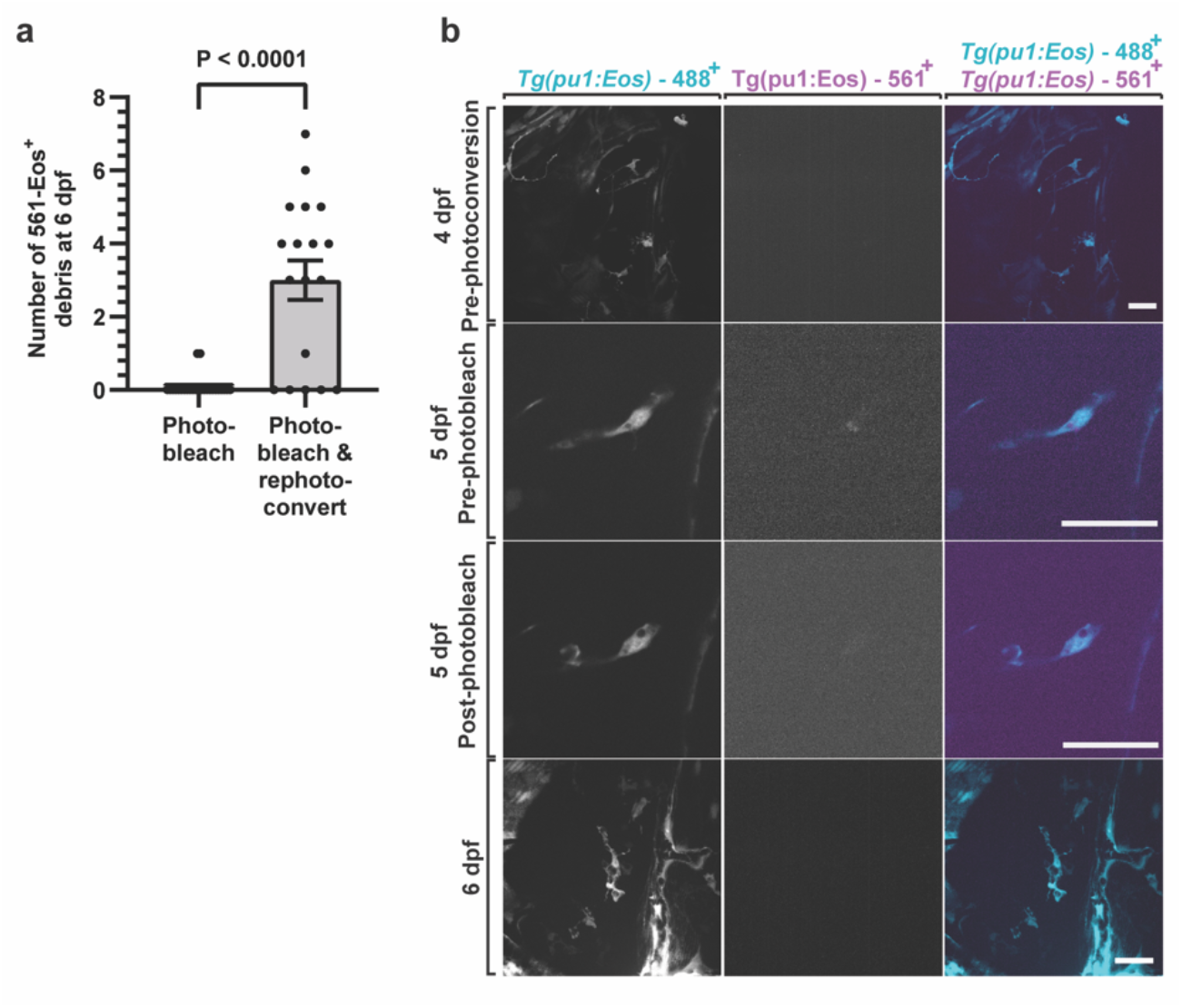
Supplemental material for Figure 3. A. Quantification of the abundance of 561-Eos^+^ debris in *Tg(pu1:Eos)* animals that had microglia photoconverted at 4 dpf, and then the debris was photobleached at 5 dpf (photo-bleach) and images were captured at 6 dpf. This is in contrast to the quantification of 561-Eos^+^ debris from animals that had microglia photoconverted at 4 dpf, photobleached at 5 dpf, rephotoconverted at 5 dpf, and then imaged at 6 dpf (p<0.0001, t-test). B. Confocal images from the experiment and quantification depicted in A. Scale bars are 25 μm (b). Descriptive statistics represented in Table S1.

**Figure S4.**
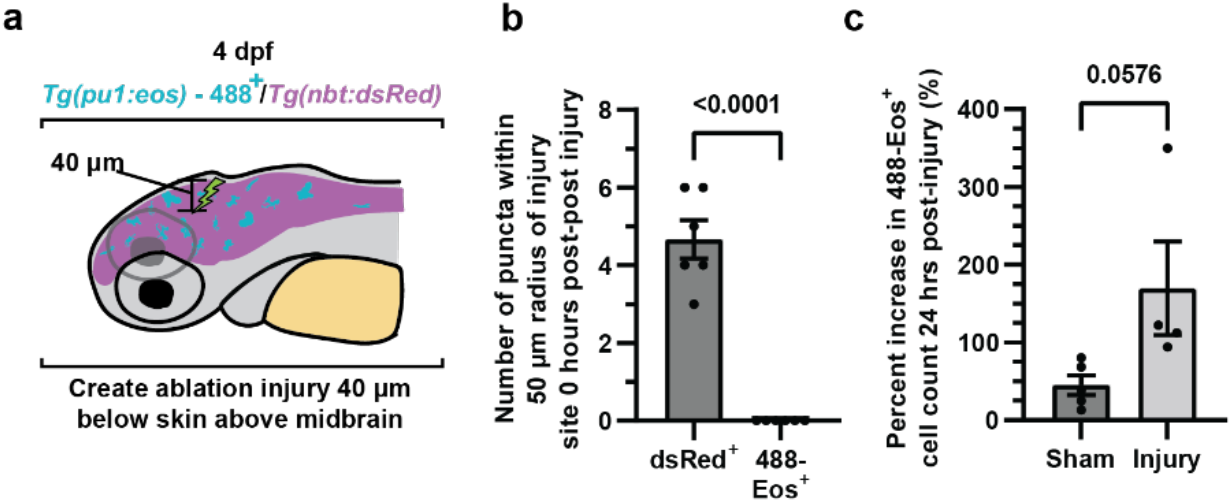
Supplemental material for Figure 4. A. Schematic demonstrating how injuries were created and calibrated in animals. B. Quantification of the amount of debris immediately after injury at 4 dpf from Tg(*nbt:DsRed)*^*+*^ vs *Tg(pu1:Eos)*^*+*^ cells in the injury paradigm (p<0.0001 dsRed^+^ vs 488-Eos^+^, unpaired t-test). C. Quantification of the percentage increase in *Tg(pu1:Eos)*^*+*^ cells in animals 24 hours after sham or injury (p=0.0576 sham vs injury, unpaired t-test). Note the microgliosis consistent with focal brain injuries. Descriptive statistics represented in Table S1.

